# Linking the dynamics of chromatin occupancy and transcription with predictive models

**DOI:** 10.1101/2020.06.28.176545

**Authors:** Trung Q. Tran, Heather K. MacAlpine, Vinay Tripuraneni, Sneha Mitra, David M. MacAlpine, Alexander J. Hartemink

## Abstract

Though the sequence of the genome within each eukaryotic cell is essentially fixed, it exists within a complex and changing chromatin state. This state is determined, in part, by the dynamic binding of proteins to the DNA. These proteins—including histones, transcription factors (TFs), and polymerases—interact with one another, the genome, and other molecules to allow the chromatin to adopt one of exceedingly many possible configurations. Understanding how changing chromatin configurations associate with transcription remains a fundamental research problem. We sought to characterize at high spatiotemporal resolution the dynamic interplay between transcription and chromatin in response to cadmium stress. While gene regulatory responses to environmental stress in yeast have been studied, how the chromatin state changes and how those changes connect to gene regulation remain unexplored. By combining MNase-seq and RNA-seq data, we found chromatin signatures of transcriptional activation and repression involving both nucleosomal and TF-sized DNA-binding factors. Using these signatures, we identified associations between chromatin dynamics and transcriptional regulation, not only for known cadmium response genes, but across the entire genome, including antisense transcripts. Those associations allowed us to develop generalizable models that can predict dynamic transcriptional responses on the basis of dynamic chromatin signatures.

## Introduction

Organisms require genic transcription to produce the proteins necessary for biological functions like growth, replication, repair, and response to environmental changes. Transcription is tightly regulated through the complex interplay of a myriad of DNA-binding factors (DBFs), including the histone octamers at the core of a nucleosome, transcription factors (TFs), and polymerases. These proteins and complexes involved in transcription, and the many others interacting with DNA, determine the chromatin landscape. How these constituents of the chromatin bind, unbind, move, and interact to regulate transcription remains an open area of research.

Numerous studies have made major strides in characterizing the roles of protein complexes involved in transcription. Chromatin immunoprecipitation (ChIP) has been used to assay binding sites of hundreds of proteins on a genomic scale, including factors involved in SAGA-dominated stress-related pathways and TFIID-dominated housekeeping pathways (Venters *et al.* 2011). Likewise, studies have probed proteins involved in the formation of the pre-initiation complex required for transcription initiation (Rhee and Pugh 2012). The role of numerous chromatin remodelers and their interactions have been characterized in detail through ChIP, proteomics, and gene expression analysis of deletion mutants (Krogan *et al.* 2006; Lenstra *et al.* 2011; Mavrich *et al.* 2008; Shivaswamy and Iyer 2008; Weiner *et al.* 2012, 2015). However, limitations in these methods, including lack of antibodies for ChIP or viability of deletion strains, are often constraining. Analysis can be complicated by the difficulty in disentangling direct chromatin effects from the pleiotropic action of the many factors and remodelers that impinge upon transcription, often indirectly. These and other issues contribute to our still limited understanding of the dynamic interplay of the chromatin landscape and transcription.

An alternative approach has been to profile chromatin occupancy in a protein-agnostic manner using nuclease digestion. Digestion by a nuclease, such as micrococcal nuclease (MNase), provides a complementary perspective to understand chromatin occupancy as it can probe accessibility at base-pair precision. Recent genome-wide mapping studies have used nucleosome-sized MNase-seq fragments to characterize the dynamics of nucleosomes under various conditions, including the cell cycle (Nocetti and Whitehouse 2016), DNA damage (Tripuraneni *et al.* 2019), and heat shock (Teves and Henikoff 2011). Additionally, studies have attempted to understand the roles of the smaller DNA-bound factors that correspond to subnucleosomal MNase-seq fragments (Belsky *et al.* 2015; Brahma and Henikoff 2019; Chereji *et al.* 2017; Henikoff *et al.* 2011; Kubik *et al.* 2017; Ramachandran *et al.* 2017; Teves and Henikoff 2011). These studies highlight the challenge of characterizing the vast heterogeneity of—and interactions among—proteins and complexes involved in DNA-mediated processes, including transcription.

Factor-agnostic chromatin occupancy profiles from MNase provide an opportunity to link changes in chromatin at nucleotide resolution with transcriptional regulation, especially regulation induced by environmental perturbations. Here, we utilize a high-resolution spatiotemporal stress response data set to elucidate the relationship between chromatin organization and gene expression by developing general strategies and models to analyze, genome-wide, chromatin dynamics relative to changes in transcription.

## Results

### Paired-end MNase-seq captures high-resolution chromatin occupancy dynamics associated with transcription during cadmium stress

We sought to characterize the dynamics of chromatin in terms of changes in occupancy and organizational structure of nucleosomes as well as smaller transcription-related proteins. A nucleotide resolution view of chromatin occupancy dynamics in response to cadmium stress would allow us to associate and infer relationships between these chromatin changes and those in transcription. Yeast cells were exposed to cadmium and samples were collected over a two-hour time course (Fig. 1A). Chromatin occupancy and positioning dynamics were profiled using paired-end MNase-seq to map DNA-binding factors at base pair resolution (Fig. 1B). Concurrently, transcripts were interrogated using strand-specific total RNA-seq (Fig. 1C).

**Figure 1.**
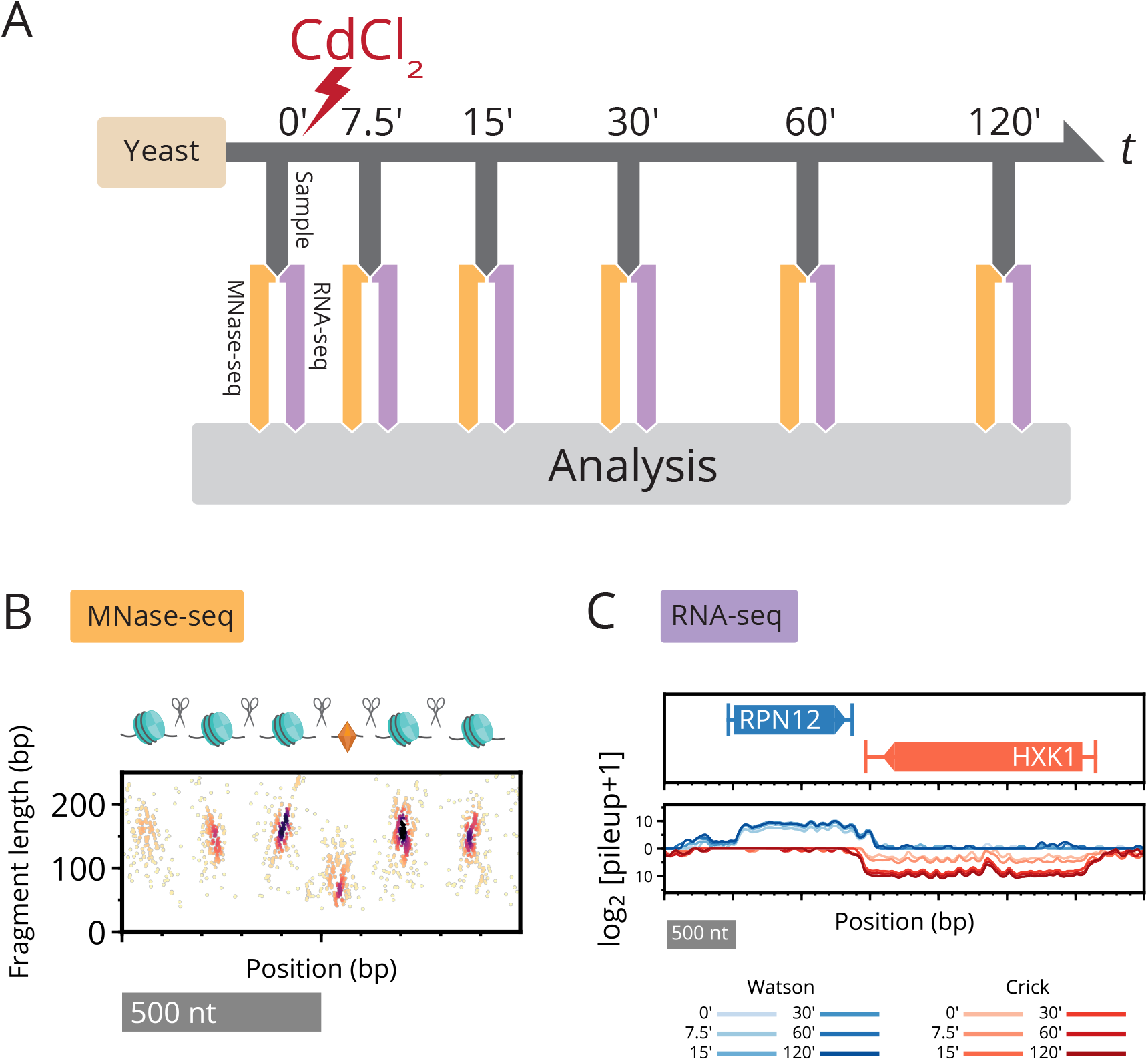
Paired-end MNase-seq and stranded RNA-seq capture high-resolution chromatin occupancy and transcriptome state throughout a perturbation time course. **(A)** Overview of cadmium perturbation experiment in which paired-end MNase-seq and strand-specific RNA-seq samples were collected immediately prior to cadmium exposure and for five additional time points over two hours. **(B)** Depiction of nucleosomes flanking a small (subnucleosomal) binding factor, and fragments that result upon digestion by MNase. Paired-end MNase-seq fragments are plotted based on their center position and length. **(C)** Strand-specific RNA-seq is plotted as the log2 pileup, the number of total RNA-seq reads at each genomic position, separately mapped to Watson (blue) and Crick (red) strands. Changing RNA-seq read levels over the time course are plotted using progressive coloring for each strand.

To evaluate our data and methods, we considered the well-studied stress response gene *HSP26*, whose role is to facilitate the disaggregation of misfolded proteins (Cashikar *et al.* 2005). Hsp26 has been implicated in responses to many stress conditions, including heat shock (Benesch *et al.* 2010; Franzmann *et al.* 2008), acidity (Kawahata *et al.* 2006), sulfur starvation (Pereira *et al.* 2008), and metal toxicity (Hosiner *et al.* 2014; Momose and Iwahashi 2001). Furthermore, several transcription factors, including Hsf1, Met4, and Met32, have been found to bind in the well-characterized promoter of *HSP26* (Boy-Marcotte *et al.* 1999; Carrillo *et al.* 2012; Chen and Pederson 1993; Susek and Lindquist 1990; Treger *et al.* 1998). Given this context, *HSP26* serves as a useful test case because we understand many aspects of its local chromatin dynamics when it is activated under stress conditions.

We observed significant changes in the chromatin around the transcription start site (TSS) of *HSP26* (Fig. 2A), coinciding with a dramatic increase in its transcript level. Upstream, in the promoter of *HSP26*, nucleosome-sized fragments of length 144–174 bp are replaced by small fragments less than 100 bp. In the gene body of *HSP26*, nucleosome-sized fragments become “fuzzy”, increasing in positional and fragment-length variability (Fig. 2A). Nucleosomes upstream of *HSP26* are known to be evicted (Lee *et al.* 2004) and replaced by smaller factors associated with transcription initiation, pushing gene body nucleosomes downstream (Fig. 2B,C). Then, active transcription by RNA polymerases displaces and evicts nucleosomes in its path (Kulaeva *et al.* 2010; Lee *et al.* 2004; Schwabish and Struhl 2004), which is apparent in our data in the significant loss of nucleosomal fragments within the gene body of *HSP26*.

**Figure 2.**
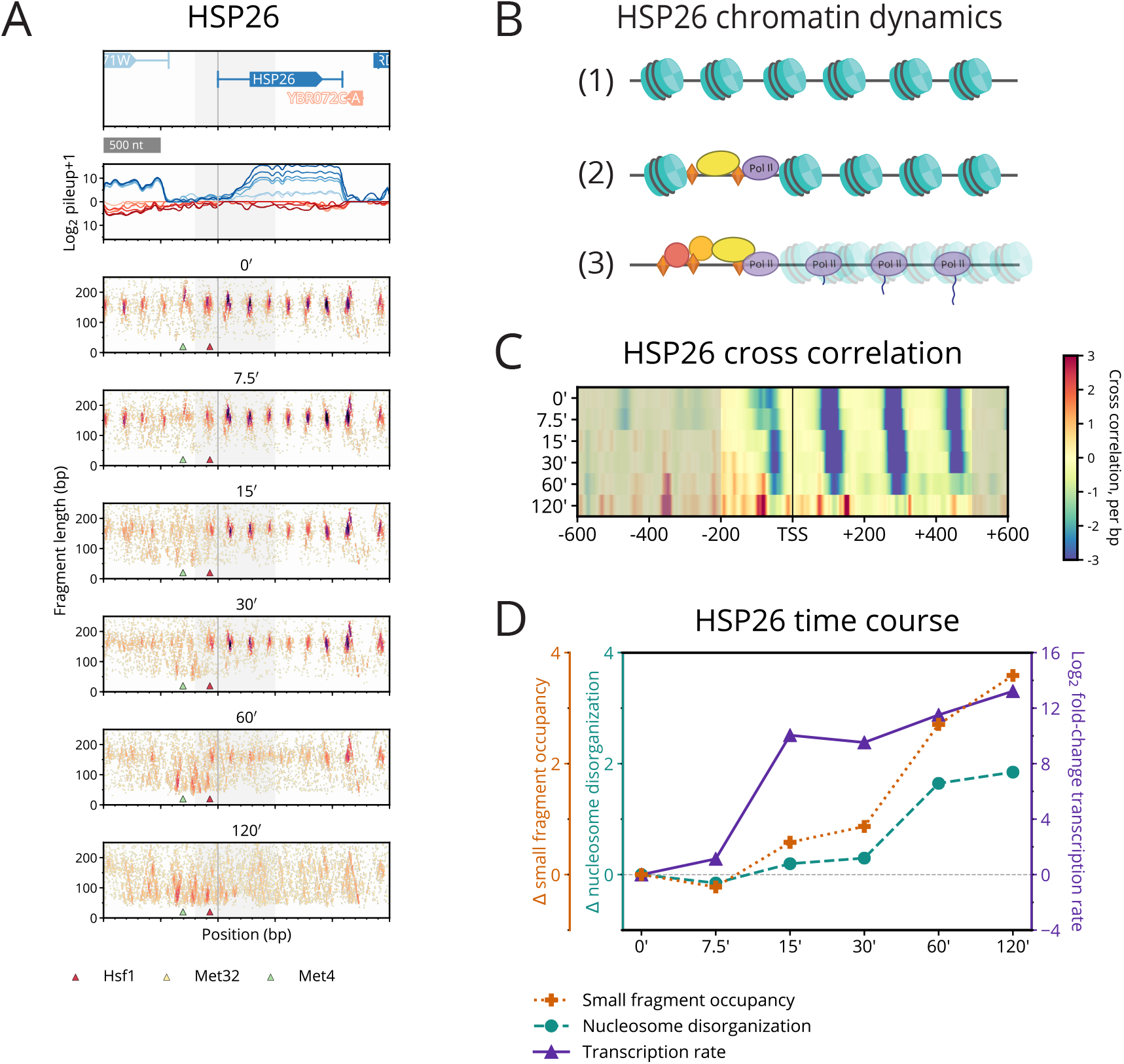
Cadmium induces local chromatin dynamics that correlate with transcription of *HSP26*. **(A)** Typhoon plot shows dynamics of MNase-seq and RNA-seq data near *HSP26*. Nucleosomes in the promoter region are replaced by small fragments, while gene body nucleosomes disorganize (grey shading highlights the [–200,500] region around the TSS that we analyze for all genes). Small fragments appear around motifs for known regulators Hsf1 (red triangle), Met4 (green triangle), and Met32 (obscured by green triangle). **(B)** Depiction of the chromatin dynamics for *HSP26*. (1) Before treatment, nucleosomes are well-positioned. (2) Between 15–30 min, nucleosomes are evicted from the promoter region and replaced by transcription-related proteins and complexes. (3) By 60–120 min, nucleosomes are fuzzy and polymerases are actively transcribing *HSP26*. **(C)** Heatmap of differential cross-correlation values of *HSP26* through the time course, summarizing how gene body nucleosomes initially shift downstream and then disappear, and how promoter nucleosomes are rapidly displaced as small fragments accumulate. Higher values (more red) indicate higher cross-correlation with subnucleosome fragments; lower values (more blue) indicate a stronger signal for nucleosome fragments. **(D)** Line plot of *HSP26* time course summarizing the change relative to 0 min in occupancy of promoter small fragments (orange), disorganization of gene body nucleosomes (turquoise), and transcription rate (purple).

To quantify these complex transcription-associated chromatin dynamics genome-wide, we defined two scores for each gene, a “small fragment occupancy” score of small fragments appearing in a gene’s promoter, and a measure of “nucleosome disorganization” within its gene body using information entropy. Additionally, to account for variations in RNA stability, we estimated transcription rates from our measured transcript levels using published mRNA decay rates (Geisberg *et al.* 2014; Miller *et al.* 2011; Presnyak *et al.* 2015).

Using these measures, we are able to succinctly describe relationships between chromatin dynamics and transcription in a range of genes, from activated *HSP26* (Fig. 2D), to repressed *RPS7A* (Supplemental Fig. 1), to unchanging *CKB1* (Supplemental Fig. 2). Averaging these two measures of the chromatin across the time course, and then ranking all genes by the resulting “combined chromatin” score, we observed large-scale coordination between chromatin and transcription across a significant proportion of the genome (Fig. 3A,B).

**Figure 3.**
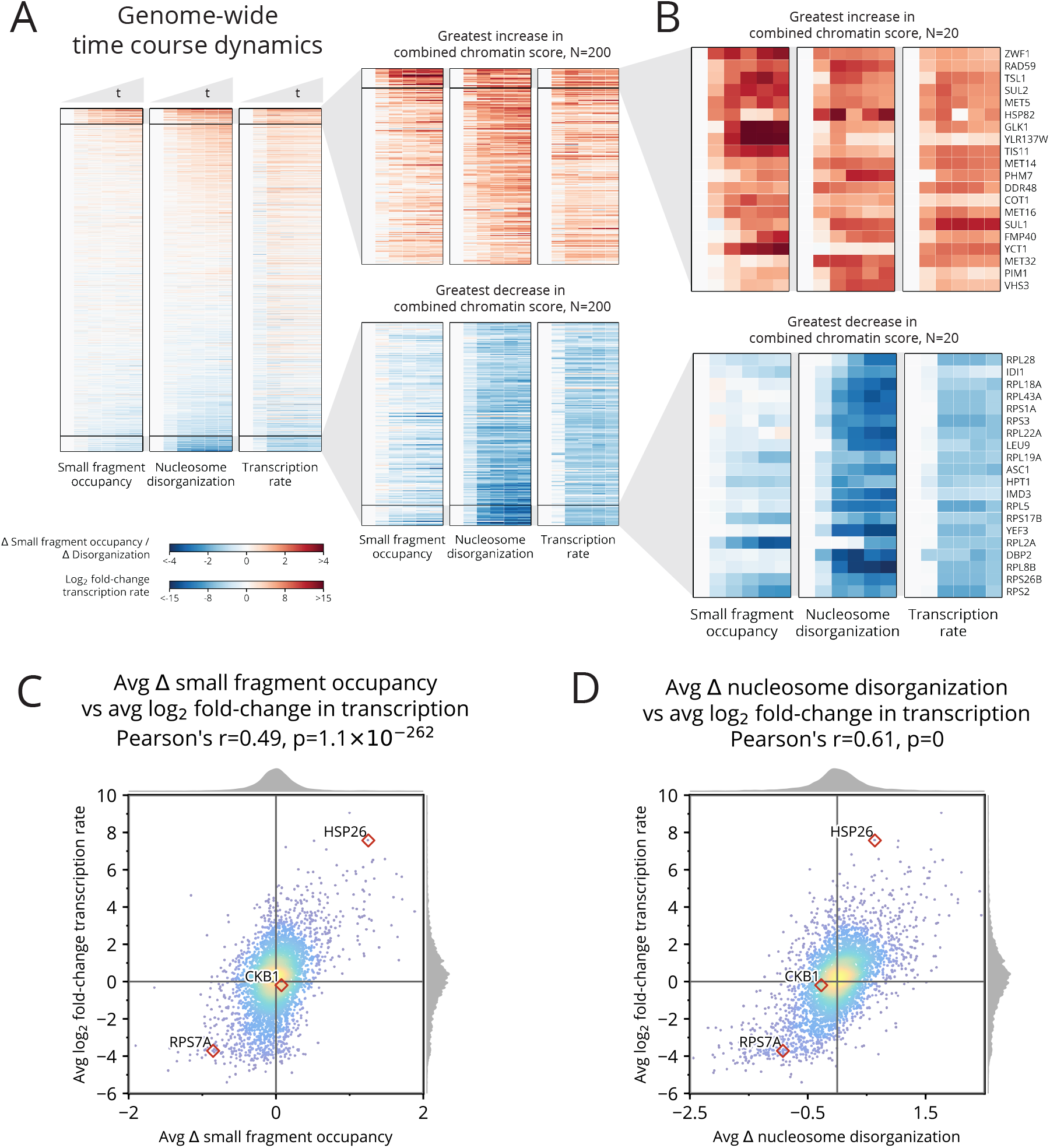
Cadmium induces genome-wide chromatin dynamics that correlate well with genome-wide transcriptional dynamics. **(A)** Heatmaps of changes in chromatin occupancy measures and transcription rate for all genes and all times, relative to 0 min (left: promoter small fragment occupancy; middle: gene body nucleosome disorganization; right: transcription rate). Genes (rows) are sorted by combined chromatin score. **(B)** Detailed heatmaps of the 20 genes whose combined chromatin scores increase (top) or decrease (bottom) most. **(C)** Scatter plot of relationship between change in small fragment occupancy and log2 fold-change in transcription rate, each averaged over the time course. **(D)** Scatter plot of relationship between change in nucleosome disorganization and log2 fold-change in transcription rate, each averaged over the time course.

Globally, log fold-changes in transcription show a significant positive Pearson correlation with changes in each of our chromatin measures: 0.49 for small fragment occupancy (Fig. 3C), 0.61 for nucleosome disorganization (Fig. 3D), and 0.68 for combined chromatin (Supplemental Fig. 3A). The high correlation between combined chromatin and transcription, along with a lower 0.33 correlation between small fragment occupancy and nucleosome disorganization (Supplemental Fig. 3B), suggests that small fragment occupancy and nucleosome disorganization each provide orthogonal statistical power in describing changes in the chromatin relative to changes in transcription.

### Changes in nucleosome and small factor occupancy at TSSs recapitulate genome-wide transcriptional response to cadmium

To determine whether chromatin dynamics alone could recapitulate known response to cadmium exposure, we performed Gene Ontology (GO) enrichment analysis of the 300 genes with the highest and lowest values for each chromatin measure. We identified, with varying levels of false discovery rate (FDR) significance, regulation pathways implicated under cadmium exposure. We further validated these chromatin-identified pathways using literature and a separate GO enrichment analysis based on changes in transcription (Supplemental Tables S1 and S2).

One of the established responses for cells undergoing stress involves shutting down ribosomal and other translation-related pathways (Hosiner *et al.* 2014; Reja *et al.* 2015; Vinayachandran *et al.* 2018). Using our simple chromatin measures, ribosomal and translation-related GO terms emerged as the most significantly down-regulated, with FDR values often much lower than 10^−10^ (Fig. 4A).

**Figure 4.**
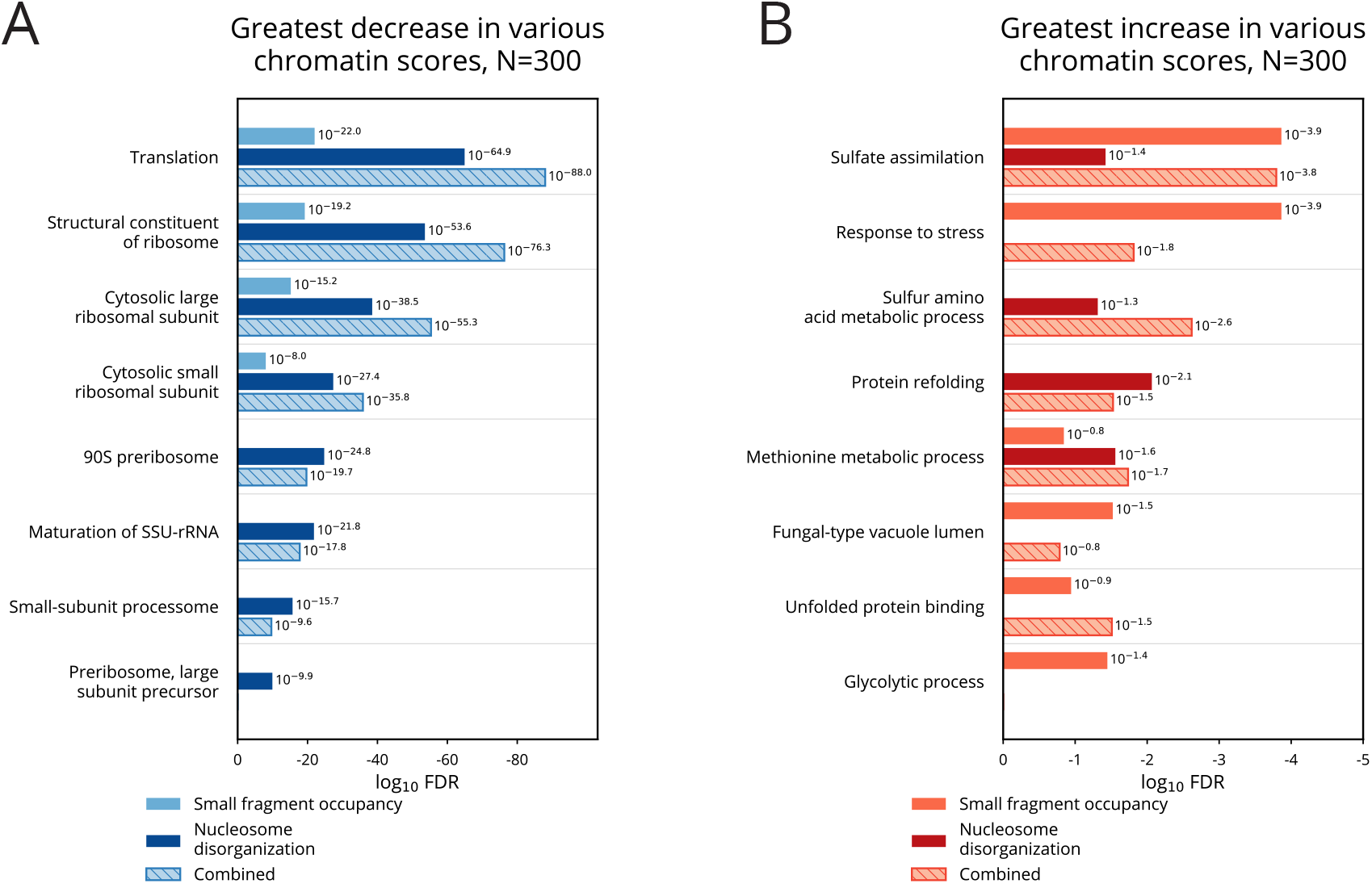
GO enrichment analysis of genes with highly dynamic chromatin recovers established cadmium response pathways. **(A)** Top 8 categories resulting from GO enrichment analysis of 300 genes with greatest decrease in small fragment occupancy, nucleosome disorganization, and combined chromatin score. Translation-related genes are recovered with significant FDR. **(B)** Top 8 categories resulting from GO enrichment analysis of 300 genes with greatest increase in small fragment occupancy, nucleosome disorganization, and combined chromatin score. Genes involved with stress response, sulfur assimilation, and protein folding pathways are recovered with significant FDR.

Translation-related genes are repressed as a tightly regulated cluster, but pathways activated under cadmium exposure are also recovered as the most significantly upregulated by our chromatin scores, albeit with FDR values above 10^−4^ (Fig. 4B). Consistent with previous cadmium and heavy metal stress response studies (Faller *et al.* 2005; Fauchon *et al.* 2002; Hartwig 2001) and our own transcriptional GO enrichment analysis (Supplemental Table S2), two major cadmium-response pathways were implicated by changes in the chromatin: sulfur assimilation and protein folding. While small fragment occupancy identified sulfate assimilation and stress response terms with the greatest significance (FDR of 10^−3.9^), nucleosome disorganization was required to identify protein refolding and sulfur amino acid metabolic process terms. Our different measures computed from chromatin are sufficient to accurately recover high-level stress response pathways induced and repressed by cadmium exposure.

### High-resolution time course recovers cascading induction of sulfur pathways

Because of the significant involvement of sulfur assimilation in the cell’s response to cadmium, we next sought to detail changes in the chromatin related to the activation of sulfur pathways. The heavy demand for sulfur arises because it is required for the biosynthesis of the cadmium-chelating glutathione (Fauchon *et al.* 2002). Sulfur pathways are activated through Met4 and its binding complex, comprised of cis-binding factors Cbf1 and Met31/Met32, and accessory factor Met28 (Blaiseau and Thomas 1998; Kuras *et al.* 1996). Met4 is negatively regulated through ubiquitination by SCF^Met30^ (Barbey *et al.* 2005; Kaiser *et al.* 2000; Kuras *et al.* 2002)(Fig. 5A). In our study, we identified novel features of the chromatin in the cascading events that regulate the sulfur metabolic pathways (Fig. 5B): (i) the activation of the Met4 complex through its cofactors, (ii) the activation of the sulfur pathways by Met4, and (iii) the subsequent down-regulation of Met4 activity by SCF^Met30^, evident in diminished transcription of Met4-regulated genes.

**Figure 5.**
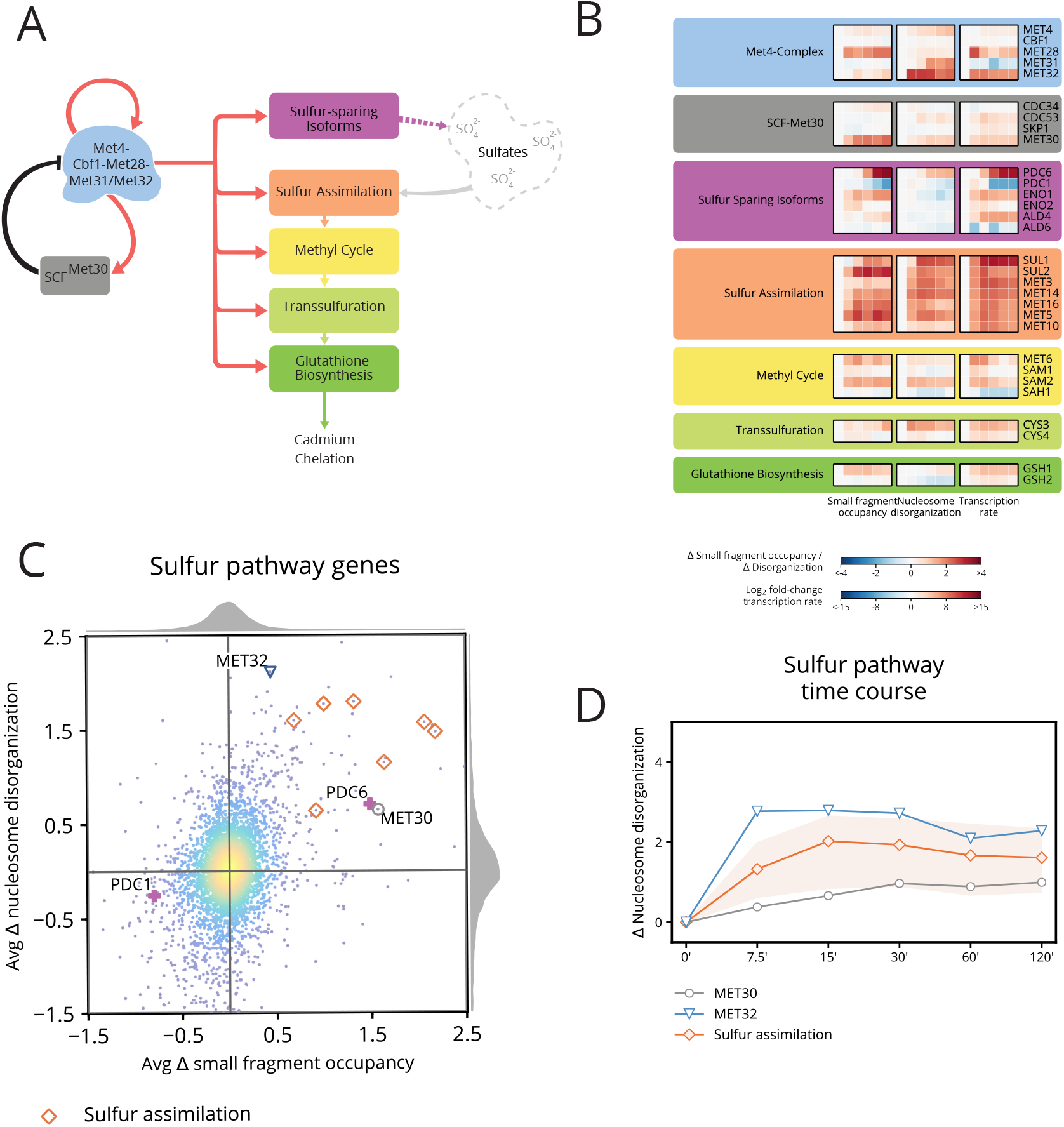
Chromatin and transcription dynamics detail Met4 and Met32 functional activation, induction of sulfur genes, and subsequent regulation. **(A)** The Met4 complex activates cascading sulfur pathways required for cadmium chelation and also activates its negative regulator SCF^Met30^. **(B)** Heatmap of changes in chromatin occupancy and transcription rate for the sulfur pathway genes. Cofactors of the Met4 complex exhibit dramatic chromatin changes in small fragment occupancy (for *MET28*) and nucleosome disorganization (for *MET32*). Sulfur sparing isoforms occur as isoenzyme pairs; members of each pair exhibit inverse chromatin dynamics (most pronounced between *PDC6* and *PDC1*). Nearly all of the sulfur assimilation pathway members show a dramatic increase in small fragment occupancy and nucleosome disorganization. **(C)** Scatter plot of average change in small fragment occupancy and average change in nucleosome disorganization. Chromatin dynamics in sulfur-related genes may manifest primarily in a single measure of the chromatin, as with *MET32* (blue triangle), *MET30* (gray circle), and *PDC6*/*PDC1* (violet), or in both small fragment occupancy and nucleosome disorganization, such as with the sulfur assimilation genes (orange diamonds). **(D)** Line plot of the change in nucleosome disorganization for the regulator gene *MET30*, activator gene *MET32*, and sulfur assimilation genes (orange line represents mean and light orange region represents full range of values across all seven genes). The disorganization for Met4 complex cofactor *MET32* is highest at 7.5 min while the sulfur assimilation genes and *MET30*, both of which are activated by the Met4 complex, reach their greatest nucleosome disorganization between 15–30 min.

Upon deubiquitination by cadmium (Barbey *et al.* 2005), Met4 becomes functionally active and induces its own cofactors (Barbey *et al.* 2005; McIsaac *et al.* 2012) and, through feedforward regulation between Met4 and Met32, activates sulfur pathway genes (Carrillo *et al.* 2012; McIsaac *et al.* 2012). We observed this activation not only in increased transcription within 7.5 minutes for *MET32* and *MET28*, but also in dramatic nucleosome disorganization of *MET32* (Supplemental Fig. 4) and increased small fragment occupancy for *MET28*.

While Met31 shares a binding motif and largely overlaps in function with Met32 (Blaiseau *et al.* 1997), it is not as prominent as Met32 in the activation of sulfur pathways (Carrillo *et al.* 2012; McIsaac *et al.* 2012; Petti *et al.* 2012). In response to cadmium, the transcription of *MET31* is repressed, but the chromatin around the gene exhibits an unexpected behavior in light of this: although *MET31* expression is repressed, its nucleosomes become highly disorganized. Leveraging our stranded RNA-seq data, we noticed significantly increasing antisense transcription over the time course (Supplemental Fig. 5). Additionally, downstream of the transcription end site (TES) of *MET31*, small fragments become enriched at a Met31/Met32 binding motif. Taken together, our data suggests that *MET31* is being regulated by non-coding RNA (ncRNA) antisense transcription.

Following activation of the Met4 complex (Carrillo *et al.* 2012; McIsaac *et al.* 2012), small fragment occupancy, nucleosome disorganization, and transcription increase for the seven sulfur assimilation genes (Fig. 5C) and many downstream genes within 15 minutes. Additionally, the Met4 complex induces a sulfur-sparing transcriptional-switch between functionally similar isoenzymes to indirectly contribute sulfur required for chelation. This switch includes replacing sulfur-rich Pdc1 with sulfur-lacking Pdc6, Ald6 with Ald4, and Eno2 with Eno1 (Fauchon *et al.* 2002). We see evidence of these substitutions between isoenzyme pairs in our data, with the most dramatic changes evident in the small fragment occupancy of *PDC6* (details in Supplemental Fig. 6) and *PDC1*.

Following induction of the sulfur pathways, the activating roles of Met32 and Met4 diminish upon regulation by SCF^Met30^ (Ouni *et al.* 2010; Patton *et al.* 2000). This regulation is observed in our data in the gradually increasing transcription and nucleosome disorganization of *MET30* throughout the time course, as well as in how the nucleosome disorganization scores of *MET32* and many of the sulfur assimilation genes gradually diminish after an early peak (Fig. 5B,D).

Together, these results and analyses complement established transcriptional and ChIP-based studies by detailing chromatin dynamics of the sulfur metabolic pathways and identifying a potentially novel regulatory mechanism for *MET31* through antisense transcription.

### Cadmium treatment induces chromatin dynamics as distinct temporal clusters, including those linked to antisense transcription

We selected the 500 genes exhibiting the greatest average increase in either small fragment occupancy or nucleosome disorganization, and performed hierarchical clustering on the resulting 832 genes (fewer than 1000 because many were in both sets). Clustering revealed distinct temporal patterns in small fragment occupancy and nucleosome disorganization among the genes (Fig. 6A). GO enrichment analysis identified different stress response pathways in two of the clusters (Fig. 6B), suggesting that chromatin changes in these pathways differ in their temporal pattern. Clusters 6–8 reveal unexpected anti-correlated relationships between chromatin and transcription for genes in these clusters. For genes in cluster 6, some of the anti-correlation can be attributed to antisense transcription (Fig. 6C), as previously highlighted in *MET31*. But in cluster 7, *MCD4*, which codes for an endoplasmic reticulum membrane protein, counter-intuitively exhibits chromatin with nucleosomes that become more organized despite increased sense and no evident antisense transcription (Supplemental Fig. 7).

**Figure 6.**
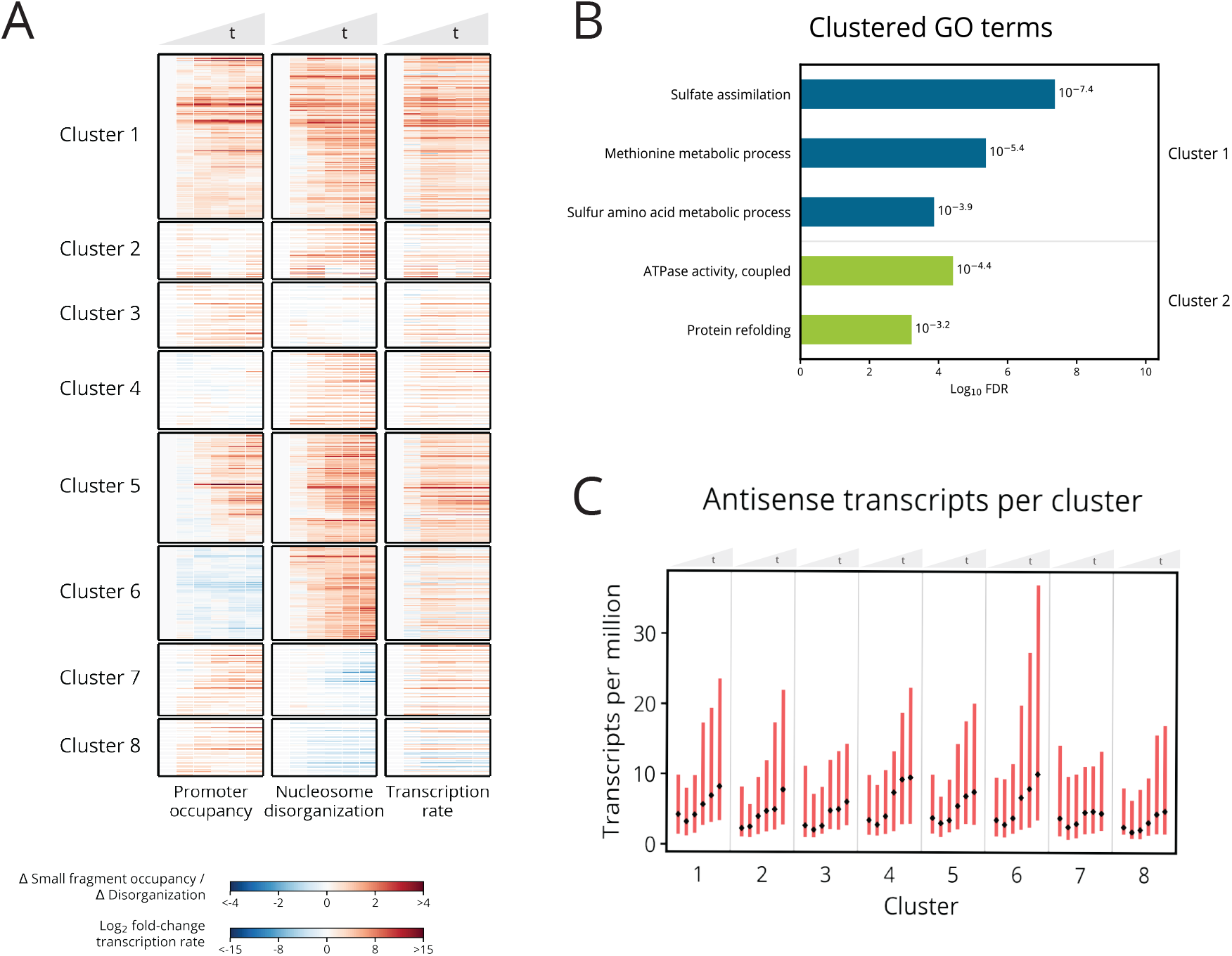
Small fragment occupancy in the promoter and gene body nucleosome disorganization reveal stress response pathway timing and patterns with antisense transcription. **(A)** Hierarchical clustering of 832 genes in the union of the 500 with greatest increase in average small fragment occupancy and the 500 with greatest increase in average nucleosome disorganization. Clusters 6–8 contain genes exhibiting anti-correlated chromatin dynamics. **(B)** GO enrichment analysis shows clusters 1 and 2 are enriched for genes in sulfur metabolism and protein refolding pathways, respectively. **(C)** Median (black dot) and interquartile range (red bar) of antisense transcript levels for genes within each cluster across the time course. Cluster 6 genes display a marked increase in antisense transcripts, perhaps explaining why the cluster exhibits increased nucleosome disorganization despite decreased small fragment occupancy in panel A.

Genome-wide, we observed that antisense transcription manifests itself with minimal apparent connection to sense transcription (Fig. 7A). Nevertheless, we did detect two general phenomena, each consistent with prior studies. First, as also seen in other environmental conditions (Kim *et al.* 2010; Till *et al.* 2018; Wilhelm *et al.* 2008), yeast undergoing cadmium stress induce pervasive antisense transcription. As the time course progresses, more and more genes exhibit increased levels of antisense transcription (Fig. 7B). Even among the 3,199 genes whose sense transcription changes only minimally, 542 exhibit at least a four-fold increase in antisense transcription (Fig. 7C). Second, previous studies have found antisense transcription can be associated with either repression or activation of target genes (Kornienko *et al.* 2013; Swamy *et al.* 2014; Till *et al.* 2018; Vance and Ponting 2014), and we observed the same phenomenon. Under cadmium stress, we identified 200 genes whose antisense transcripts increased at least four-fold and whose sense transcripts changed by at least four-fold. Among those, 104 had repressed sense transcription—*e.g.*, *MET31* and *UTR2*, whose overexpression has been linked with endoplasmic reticulum stress (Miller *et al.* 2010) (Supplemental Fig. 8)—but 96 had activated sense transcription, including the gene *YBR241C* (Supplemental Fig. 9), coding for a vacuole localization protein (Wiederhold *et al.* 2009).

**Figure 7.**
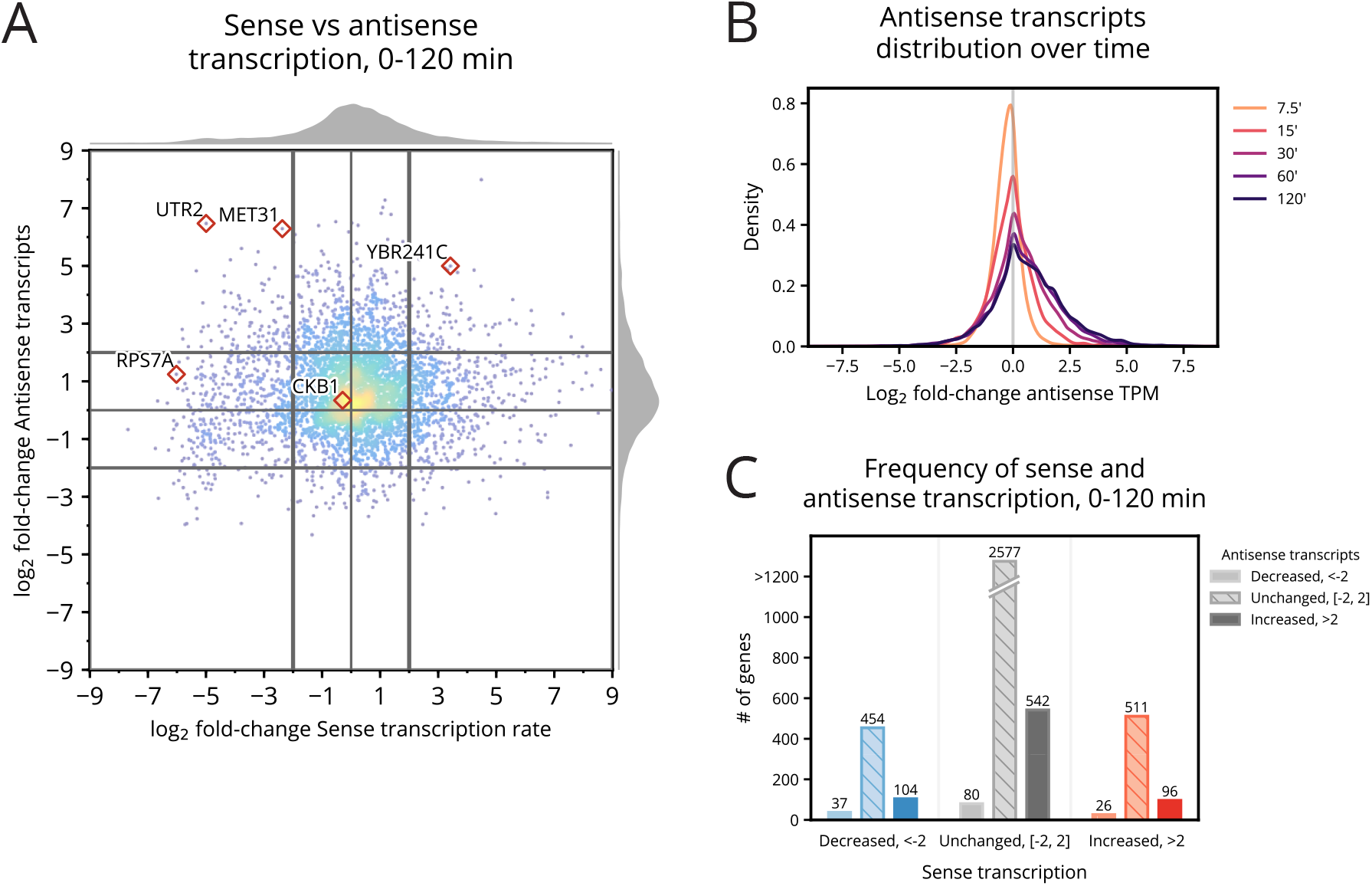
Cadmium induces changes in both sense and antisense transcription. **(A)** Distribution of the log2 fold-change in sense transcription against the log2 fold-change in antisense transcripts from 0–120 min. Antisense transcripts are enriched genome-wide by 120 min. **(B)** Distribution of the log2 fold-change in antisense transcripts for each time point following 0 min. Antisense transcripts monotonically increase throughout the time course. **(C)** Counts of genes that exhibit decreased, unchanged, and increased sense and antisense transcripts from 0–120 min. Genes in each category of sense transcription exhibit positively skewed enrichment of antisense transcripts.

**Figure 8.**
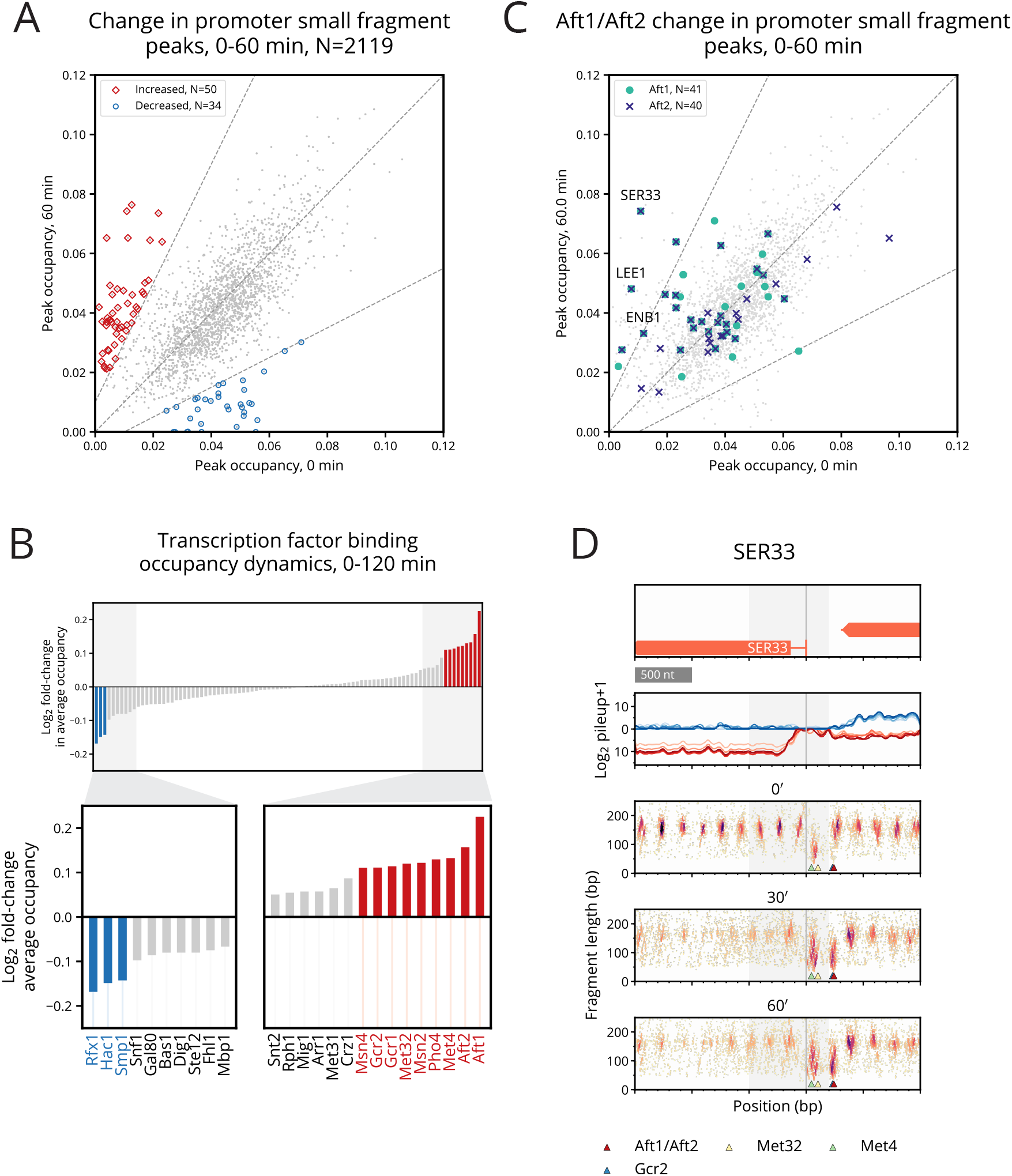
Small fragment occupancy in the promoter reveals transcription factor binding dynamics implicated in cadmium stress response. **(A)** Scatter plot of the 0–60 min occupancy change for 2,119 small fragment peaks identified in gene promoters. 50 peaks increased in occupancy by at least double (red), while 34 peaks decreased by at least half (blue). **(B)** Average change in occupancy for promoter peaks per FIMO-assigned TF. TFs are labeled as increased/decreased (red/blue) if the absolute value of their average log-fold change exceeds 0.1. TFs with the greatest increase in binding occupancy include the iron homeostasis regulators Aft1/Aft2, sulfur pathway regulators Met4/Met32, glycolytic activators Gcr1/Gcr2, and general stress responders Msn2/Msn4. **(C)** Scatter plot of occupancy of Aft1 (turquoise circle) and Aft2 (blue X) at 0 min and 60 min. Aft1 and Aft2 exhibit genome-wide enrichment in binding at 60 min compared to 0 min, particularly in the promoters of a few genes like *SER33*. **(D)** Typhoon plot of *SER33* shows small fragment enrichment at Aft1/Aft2 (red triangle) and Gcr2 (blue triangle, mostly obscured by red triangle) motifs as well as near Met32 (yellow triangle) and Met4 (green triangle) motifs.

### Motif analysis identifies per-bp binding dynamics of transcription factors

To explore small fragment occupancy more closely, we identified peaks in the signal and quantified the change in binding at each peak over 60 minutes (Fig. 8A). We ran the motif finder FIMO (Grant *et al.* 2011) near peak locations to associate peaks with TFs, and then for each TF, computed its average change in binding occupancy (Fig. 8B). TFs exhibiting the greatest average increase in occupancy include not only the sulfur pathway activators Met4 and Met32, general stress regulators Msn2 and Msn4, and glycolytic activators Gcr1 and Gcr2, but also the iron homeostasis regulators Aft1 and Aft2. Genes with the greatest increase in both Aft1 and Aft2 binding include *SER33*, *LEE1*, and *ENB1*.

For *SER33*, a gene involved in Ser and Gly biosynthesis (Albers *et al.* 2003), we see evidence of Aft1/Aft2 binding near Gcr2 in the promoter (Fig. 8C). Whereas Gcr2 is known to interact with Gcr1, a known regulator of *SER33* (Hu *et al.* 2007), Aft1 and Aft2 have yet to be identified as regulators for *SER33* (Fig. 8D). Additionally, we see enrichment of small fragments near the motifs for known regulators Met32 and Met4, previously identified through ChIP (Carrillo *et al.* 2012). Similarly strong evidence of Aft1/Aft2 binding is found in the promoters of *LEE1* (Supplemental Fig. 10A), a zinc-finger of unknown function, and *ENB1* (Supplemental Fig. 10B), a ferric enterobactin transmembrane transporter (Heymann *et al.* 2000). *ENB1* has only been identified to be regulated by Aft1 through microarrays (Hu *et al.* 2007). While the iron homeostasis pathways have been previously implicated in heavy metal stress conditions (Halimaa *et al.* 2019; Hosiner *et al.* 2014), our analysis further elucidates the binding dynamics of regulators Aft1 and Aft2 under cadmium stress and, more generally, demonstrates the richness of small fragment signals in MNase-seq data.

### Chromatin occupancy changes are predictive of changes in gene expression

Finally, we sought to develop a model to quantify the relationship between our measures of chromatin dynamics and changes in transcription. We used Gaussian process regression models to predict the transcription at each time point based solely on chromatin dynamics and initial transcript levels (at 0 min, before cadmium treatment). We constructed four models to evaluate the inclusion of various measures of the chromatin, culminating in a “full” model that incorporates additional occupancy measures, nucleosome positional shifts (Supplemental Fig. 11), and chromatin measures relative to called antisense transcripts (Supplemental Fig. 12).

Under 10-fold cross-validation, we evaluated each model using the coefficient of determination (*R*^2^), as the proportion of variance each model is able to explain (Fig. 9). For each feature-containing model, prediction performance gradually worsens through the time course as genes’ transcript levels increasingly diverge from their initial values. However, models that include chromatin features consistently outperform a model that just uses initial transcript levels (RNA only), with the gap growing over time. Nucleosome disorganization is more informative than small fragment occupancy, especially at intermediate times; consistent with our other results, combining both measures provides more predictive power than either alone. The full model does not add much to this combination at 7.5 and 15 minutes because early predictions are mainly driven by initial transcript levels. However, by 30 minutes, it begins to outperform all other models, maintaining an *R*^2^ of 0.44 even two hours after the cell’s exposure to cadmium.

**Figure 9.**
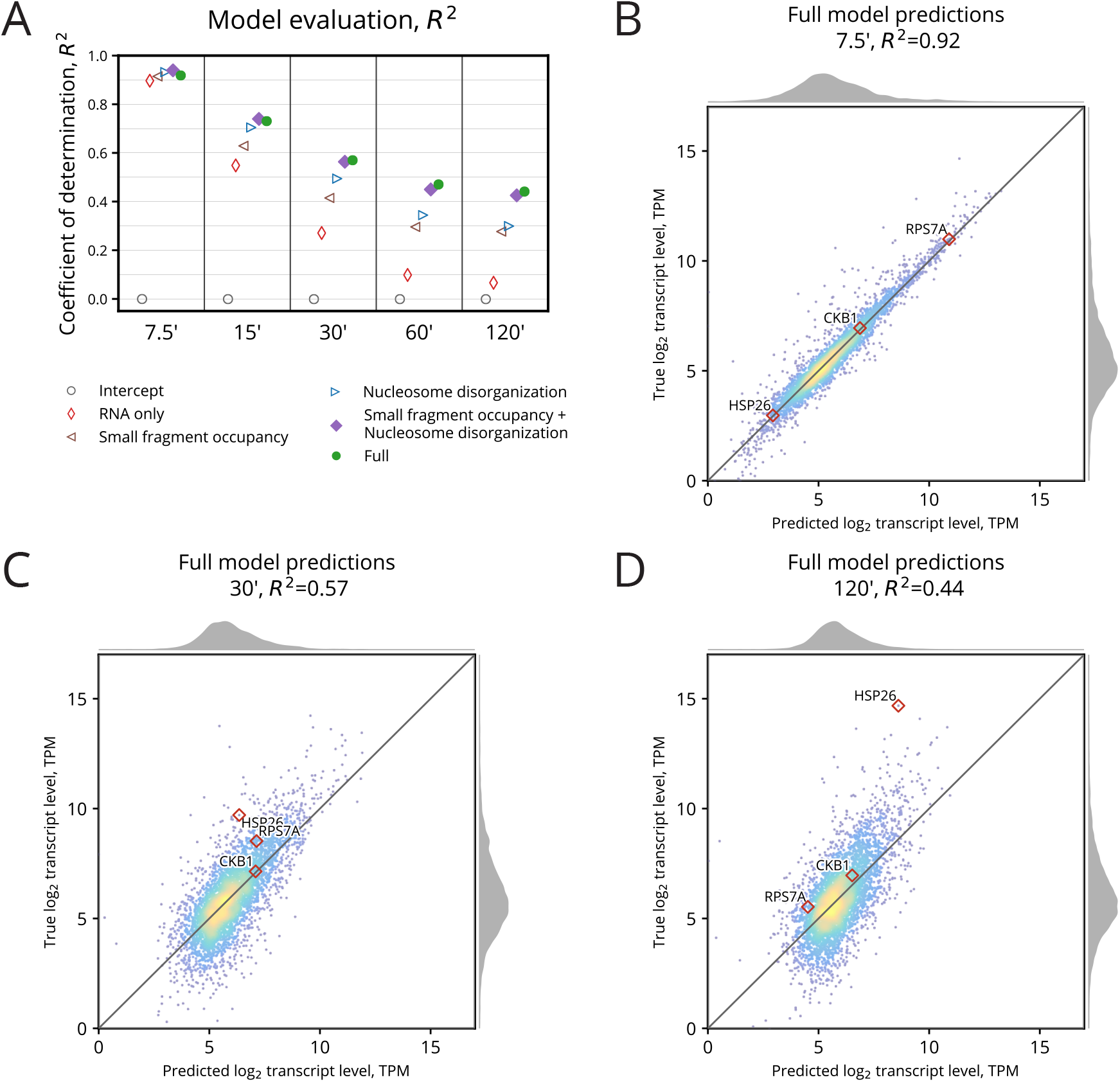
Chromatin occupancy dynamics are predictive of gene expression. **(A)** Comparison of each GP model’s performance using its coefficient of determination, *R*^2^. The Full model incorporating all chromatin features and 0 min transcript level outperforms all other models for 30–120 min. Later time points rely less on 0 min transcript level for prediction, so the marginal gain in statistical power between features becomes more evident. **(B)** Comparison between true and predicted log2 transcript level for the Full model after 7.5 min. Most genes are well predicted using 0 min transcript level. **(C)** Full model predictions at 30 min. Predictions remain well correlated, but less than at 7.5 min. **(D)** Full model predictions at 120 min. After two full hours have elapsed, transcript level predictions have become a bit less correlated, but still, *R*^2^ remains 0.44.

While our models cannot ascertain causal links between changes in chromatin and transcription and use measures that do not fully characterize the chromatin state, they nevertheless provide strong evidence that a large proportion of a cell’s transcription state can be predicted from simple measures of its chromatin state, even after significant environmental perturbation.

## Discussion

In contrast to ChIP-based studies that profile one DNA-binding factor at a time, our study surveys the occupancy of all factors across the entire genome, albeit without explicit information on their identities. While nucleosomes are well-characterized by MNase digestion, profiling the TFs and complexes that regulate gene expression is a more challenging, open problem. Prior work has explored the dynamics of various individual promoter-binding factors including TFs, general TFs, polymerases, mediator, SAGA, TFIID, chromatin remodelers and histone modifications, and others (Chereji *et al.* 2017; Huisinga and Pugh 2004; Reja *et al.* 2015; Rhee and Pugh 2012; Shivaswamy and Iyer 2008; Venters *et al.* 2011; Vinayachandran *et al.* 2018; Weiner *et al.* 2012, 2015). These studies—along with motif analysis—provide us with useful context to understand the dynamics of small fragments in our MNase-seq data, such as in our characterizations of *HSP26* (Fig. 2), the sulfur pathways (Fig. 5), and the iron homeostasis regulators Aft1/Aft2 (Fig. 8).

Analysis of *MET31*, encoding a Met4 cofactor, revealed chromatin changes linked with increased antisense transcription that may explain how the cell regulates its sense transcription. Moreover, we observed pervasive antisense transcription under cadmium stress, and while this has previously been shown to occur under a variety of environmental perturbations (Camblong *et al.* 2007; Nadal-Ribelles *et al.* 2014; Swamy *et al.* 2014; Toesca *et al.* 2011), we were able to characterize relationships between sense and antisense transcription with regulatory insight from the perspective of the local chromatin landscape. For the 667 genes we identified with antisense transcripts, including chromatin measures relative to those transcripts improved our model (marginally) for predicting sense transcription (Fig. 9A). This benefit can be further explored by narrowing in on the effect size of these antisense-related chromatin measures and by examining the individual sets of genes whose gene expression appears to have a relationship with antisense transcription.

Using just the initial transcript level and simple measures of chromatin dynamics, our regression model is able to predict the level of sense transcript with an *R*^2^ at least 0.44, even two hours after cadmium exposure (Fig. 9A). This model can be extended in multiple directions. We can further quantify the chromatin by including additional classes of fragments, by computing new measures of chromatin dynamics, and by considering chromatin beyond 200 bp of a promoter and the first 500 bp of a gene body. Additionally, our data could be modeled with other statistical methods including generalized linear models, deep neural networks, or random forests. This model and its predictions serve as a baseline showing the potential modeling opportunities and richness of statistical power of MNase-derived time-series chromatin data.

## Materials and Methods

### Yeast strain

The yeast strain used in this study has the W303 background with the genotype: MATa, leu2-3,112, trp1-1, can1-100, ura3-1, ade2-1, his3-11,15.

### Growing and sampling cells over the time course

Cells were grown asynchronously in YEPD at 30°C to an OD_600_ of 0.8. Immediately before the addition of CdCl_2_, one sample was removed and cross-linked with formaldehyde to a final concentration of 1% for MNase-seq, and another was pelleted and flash frozen for RNA-seq; these represent time 0. After the addition of CdCl_2_ to a final concentration of 1 mM, samples were taken at 7.5 min, 15 min, 30 min, 60 min, and 120 min, and processed in the above manner, respectively, in preparation for MNase-seq and RNA-seq. All experiments were repeated independently as biological replicates.

### Preparing chromatin

Cells were resuspended with 20 ml of buffer Z (0.56 M sorbitol, 50 mM Tris at pH 7.4) and 14 *μ*L of *β*-ME and 0.5 ml of a 10 mg/mL solution of zymolyase (Sunrise Science Products) prepared in buffer Z were added. Samples were incubated for 30 min at 24°C with shaking. Cells were centrifuged at 1500 rpm for 6 min at 4°C and then resuspended in 2.5 ml of NP buffer (1 M sorbitol, 50 mM NaCl, 10 mM Tris at pH 7.4, 5 mM MgCl_2_, 1 mM CaCl_2_), supplemented with 0.5 mM spermidine, 0.007% *β*-ME, and 0.075% NP-40. To determine the best digestion conditions, a four-step titration of 15 U/*μ*L MNase (Worthington) was added to 400 *μ*L of zymolyase treated cells. Samples were inverted to mix and digested on the benchtop for 20 min. The reaction was halted by adding 100 *μ*L of stop buffer (5% SDS, 50 mM EDTA). Next, proteinase K was added to a 0.2 mg/mL final concentration, and the samples were inverted and then incubated overnight at 65°C. DNA was recovered by phenol/chloroform extraction and isopropanol precipitation.

### Preparing MNase sequencing libraries

Illumina sequencing libraries of MNase-treated DNA were prepared using 500 ng of DNA as previously described (Henikoff *et al.* 2011).

### Preparing RNA sequencing libraries

Illumina sequencing libraries of total RNA were prepared using the Illumina TruSeq Stranded Total RNA Human/Mouse/Rat kit (Cat number RS-122-2201) following the protocol provided by Illumina with Ribo-Zero.

### Aligning sequencing reads to the genome

All reads were aligned to the sacCer3/R64 version of the *S. cerevisiae* genome using Bowtie 0.12.7 (Langmead *et al.* 2009).

The recovered sequences from all paired-end MNase reads were truncated to 20 bp and aligned in paired-end mode using the following Bowtie parameters: `--wrapper basic-0 --time -p 32 -n 2 -l 20 --phred33-quals -m 1 --best --strata -S`.

The recovered sequences from all single-end RNA reads were truncated to 51 bp and aligned in single-end mode using the same Bowtie parameters.

### Processing reads from MNase-seq and RNA-seq replicates

After confirming high concordance between them, MNase-seq and RNA-seq replicates were subsampled and merged to increase read depth and reduce bias from library preparation, sequencing, and digestion. Details of the procedure used to subsample and merge each pair of replicates are provided in Supplemental Method S1. After merging, we had 24,152,389 mapped MNase fragments (pairs of reads) and 42,107,377 mapped RNA reads for each time point.

### Selecting a set of genes for analysis

We compiled a set of 4,427 genes for analysis. A gene was chosen if it satisfied five criteria: it (i) is classified as either verified or uncharacterized by sacCer3/R64, (ii) contains an open reading frame (ORF) at least 500 bp long, (iii) contains an annotated TSS, (iv) has a reported mRNA half-life, and (v) has adequate MNase-seq coverage.

Genes whose ORFs are less than 500 bp (Supplemental Fig. 13A) long were omitted in order to ensure valid “gene body” calculations between [TSS, +500]. TSS annotations were determined by Park *et al.* (2014). For five genes, *SUL1*, *SUL2*, *MET32*, *HSP26*, and *BDS1*, we manually annotated the TSS to be consistent with the RNA-seq data in this study. We required a half-life for each gene in order to estimate transcription rates. MNase-seq coverage was computed in a 2,000 bp window centered on each gene’s TSS. A position in this window is considered “covered” when there exists at least one fragment whose center is at this position. MNase coverage was then defined as the number of covered positions in this window divided by the length of the window, 2,000 bp. Genes with MNase coverage below 0.85 (*n*=109) were excluded from further analysis (Supplemental Fig. 13B).

### Defining classes of MNase-seq fragments and measures of their occupancy

MNase-seq fragments can be associated with different DNA-binding factors of the basis of their length (Supplemental Fig. 14). To summarize the chromatin occupancy of different factors around genes, fragments were first filtered into two classes: fragments associated with nucleosomes, those between 144–174 bp long, and fragments associated with smaller factors, those less than 100 bp long. In determining these lengths, we made use of two reference data sets, as described next.

Nucleosomal fragment lengths were determined by examining the distribution of MNase-seq fragments prior to cadmium treatment around the top 2,500 unique nucleosome positions reported by a highly sensitive chemical assay (Brogaard *et al.* 2012). In our MNase-seq data, the distribution of fragment lengths at these sites had a clear mode at 159 bp; we chose a ±15 bp interval around this mode to capture most of the nucleosomal fragments, resulting in the final 144–174 bp range.

As for fragments associated with smaller factors, because prior studies have found clear enrichment of small fragments at Abf1 sites (Henikoff *et al.* 2011), we examined the distribution of our fragments prior to cadmium treatment around 279 Abf1 binding sites, as determined by phylogenetic conservation and motif discovery, obtained from http://fraenkel-nsf.csbi.mit.edu/improved_map/p001_c2.gff (MacIsaac *et al.* 2006). In our MNase-seq data, most of the fragments at these sites were shorter than 100 bp (mode: 75 bp), so those were classified as small fragments.

For each gene, two regions were defined relative to its TSS. The promoter region was defined as a 200 bp region upstream of the TSS, [–200, TSS]. The length of this region was chosen as previously described (Lubliner *et al.* 2013; Smale and Kadonaga 2003). The gene body region was defined as a 500 bp region downstream of the TSS, [TSS, +500], to include the +1, +2, and +3 nucleosomes.

The occupancy of a class of fragments within a particular region is computed simply as the number of fragments of that class whose centers lie within that region.

### Computing chromatin scores with cross-correlation kernels

Some chromatin statistics require more spatial precision than occupancy provides, for example when determining a factor’s position or organization. In these cases, we used cross-correlation scores in a similar manner to that described in Tripuraneni *et al.* (2019). Around each gene’s TSS, a per-bp cross-correlation score was computed to smooth positional variation and filter out non-relevant fragments. We constructed three two-dimensional cross-correlation kernels: an idealized, well-positioned nucleosome kernel (Supplemental Fig. 15A), a clearly bound small factor kernel (Supplemental Fig. 15B), and a triple-nucleosome gene body summary kernel (Supplemental Fig. 15C). Each kernel was applied to the region local to each gene’s TSS for each time point to compute a per-bp cross-correlation score (Supplemental Fig. 15D).

The nucleosome and small factor kernels were constructed using a bivariate Gaussian distribution parameterized by the mean and variance for the position and length for MNase-seq fragments. The parameters for each kernel were determined using the fragment length and position distributions at positions in Brogaard *et al.* (2012) and MacIsaac *et al.* (2006), as described in the previous subsection.

To summarize the gene body chromatin as a whole, a triple nucleosome kernel was constructed to dampen the effect of the +1 nucleosome becoming more poised to be well-positioned (Mavrich *et al.* 2008; Nocetti and Whitehouse 2016). The triple nucleosome kernel was constructed by repeating the nucleosome kernel and increasing the variance to take into account variable linker spacing. The nucleosome kernel spacing was determined using the average peak spacing between the [+1,+2] and the [+2,+3] nucleosome cross-correlation scores (Supplemental Fig. 15E).

### Quantifying nucleosome disorganization

For each gene, a random variable *X* was defined with *n* possible outcomes representing each position to evaluate relative to the gene TSS.

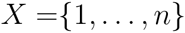

The probability of each outcome is estimated using the triple nucleosome cross-correlation scores previously defined and normalized to sum to 1.

Because the triple kernel computes a score for three approximately adjacent nucleosome positions, we set *n* = 150 to summarize the disorganization of the first three nucleosomes in the gene body starting with +1 within the [0, 150] window.

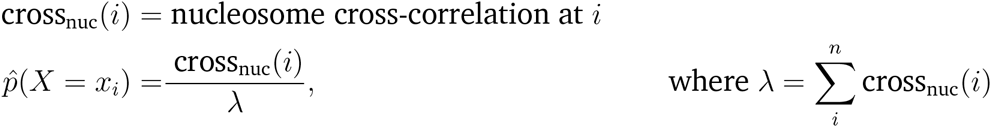

Using this random variable, a score was computed for each gene to define its “nucleosome disorganization” using information entropy (Supplemental Fig. 15F):

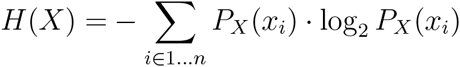

### Calling +1, +2, +3 nucleosomes and linking them over time

Nucleosomes were called using peaks of the nucleosome cross-correlation scores local to each gene’s TSS. Peaks within a 1000 bp window around the TSS were sorted by score. The position with the greatest peak score was labeled as a nucleosome center and removed. Positions within 80 bp were also removed. This procedure was repeated until all peak positions were removed and nucleosomes called for this 1000 bp window.

“Linked” nucleosomes are defined as nucleosomes across the time course that nominally represent the same underlying nucleosome even though its position may have shifted or become more or less fuzzy. Nucleosomes were linked across time points using a nearest-neighbor approach. In a greedy manner, the most well-positioned nucleosome (lowest disorganization score) was considered first. The position of this nucleosome was used to identify the linked nucleosomes in previous and subsequent time points by considering the nearest nucleosome in each of the respective time points within 100 bp of the original nucleosome’s position.

Each gene’s +1 nucleosome was called by identifying the linked nucleosome closest to the TSS. The +2 and +3 nucleosomes were computed as the next nucleosomes at least 80 bp downstream from the preceding one.

### Analyzing Gene Ontology enrichment

GO enrichment analysis was performed using GOATOOLS (Klopfenstein *et al.* 2018) with the go-basic.obo annotations from the Gene Ontology Consortium (Ashburner *et al.* 2000; The Gene Ontology Consortium 2019). False discovery rate was corrected using the Benjamini-Hochberg procedure (Benjamini and Hochberg 1995).

### Identifying transcription factor binding sites

TF binding sites were called using the small fragment cross-correlation scores in each gene promoter. The cross-correlation scores at each position and time point were sorted by score. The position with the greatest score was removed and labeled as a small fragment peak. Positions within 50 bp of the peak at any time point were also removed. This procedure was repeated until all positions were removed for each gene promoter. A small fragment occupancy value 100 bp around each peak was computed at each time point to identify positions with the greatest change in binding.

The motif finder FIMO (Grant *et al.* 2011) was run against each called peak position against the motif database from MacIsaac *et al.* (2006) using the default p-value threshold. Selected binding sites with supporting literature were annotated on typhoon plots.

### Estimating transcription rates

As previously described in Cashikar *et al.* (2005); Rabani *et al.* (2011); Yang *et al.* (2003), transcription rates were computed by incorporating mRNA decay rates into difference equations describing zero-order growth with first-order decay. Details of the procedure used to compute these transcription rates are provided in Supplemental Method S2.

### Clustering genes based on chromatin measures

Genes with the greatest increase in average small fragment occupancy or average nucleosome disorganization were chosen for clustering. The top 500 genes for each measure were combined into a final set of 832 (fewer than 1000 because many genes were in both sets).

Clustering was performed in SciPy (Virtanen *et al.* 2020) using hierarchical clustering on the basis of pair-wise Euclidean distance between z-normalized measures of change in small fragment occupancy and nucleosome disorganization. Ward linkage was chosen for its efficient approximation to the minimal sum of squares objective (Ward 1963). Eight clusters were ultimately chosen to balance the interpretability of fewer clusters with the significance of identified GO terms in smaller and more homogeneous but more numerous clusters.

### Identifying and quantifying antisense transcripts

TSSs and transcription end sites (TESs) for antisense transcripts were determined using RNA-seq pileup, the number of reads covering a genomic position. To increase signal and decrease noise, at each genomic position we added the antisense pileup values across time points to produce a cumulative pileup, and then smoothed that with a Gaussian kernel.

Starting with the highest cumulative pileup value within a gene’s transcript boundary on the antisense strand, the antisense TSS and TES were identified by progressively searching upstream and downstream, respectively, to identify the positions at which the cumulative pileup values were minimized (Supplemental Fig. 12A). Antisense transcripts were not called if they did not meet a minimum threshold of pileup at any position within the transcript boundary.

For the 667 genes where an antisense transcript could be called (Supplemental Fig. 12B), antisense transcription levels were quantified using a TPM calculation (Wagner *et al.* 2012) for strand-specific RNA-seq reads on the antisense strand within the respective antisense transcript boundaries. We also computed nucleosome disorganization and promoter occupancy chromatin measures relative to these called antisense transcripts, as previously described for the sense strand.

### Predicting transcript levels using Gaussian process regression models

Gaussian process regression models were constructed to predict the log2 transcript level for each time point using the log2 transcript level and features of the chromatin at 0 minutes, along with features of the chromatin for the time being predicted.

Four models were constructed to compare various combinations of measures of the chromatin: a small fragments promoter occupancy model, a gene body nucleosome disorganization model, a combined chromatin model, and a full model incorporating all previous models’ features with the addition of nucleosome occupancy within the promoter and within the gene body, small fragment occupancy within the gene body, +1, +2, and +3 nucleosome position shift relative to 0 min (Supplemental Fig. 11), and measures of chromatin relative to called antisense transcripts (Supplemental Fig. 12).

Each Gaussian process regression model developed using scikit-learn (Pedregosa *et al.* 2011) with a radial-basis function (RBF) kernel with length scale bounded between 0.1 and 100 and a white kernel with noise level 10^−4^ as priors for covariance. The length scale bounds and noise parameters were determined empirically through a sensitivity analysis on a subset of the data.

Promoter occupancy and nucleosome disorganization measures were each log transformed to yield an approximately normal distribution. Then, each chromatin measure (including nucleosome shift) was z-normalized to allow the RBF length parameter to be successfully approximated.

Performance for each model was evaluated using the coefficient of determination, *R*^2^, under 10-fold cross-validation.

## Data Accession

All raw and processed sequencing data generated in this study have been submitted to the NCBI Gene Expression Omnibus (GEO; https://www.ncbi.nlm.nih.gov/geo/) under accession number GSE153609.

Code to reproduce the results in this study is included in Supplemental Code and available on GitHub (https://github.com/HarteminkLab/cadmium-paper).

## Acknowledgments

The authors would like to thank Dr. Yulong Li and Dr. Greg E. Crawford for their suggestions and helpful comments during the development of this research. The work was supported by the following grants from the NIH National Institute of General Medical Sciences: R35-GM127062 (D.M.M.) and R01-GM118551 (A.J.H.).

