## Supplemental Materials for "Linking the dynamics of chromatin occupancy and transcription with predictive models"

### Contents


| | |
|---|---|
| Supplemental Figure S1 | 2 |
| Supplemental Figure S2 | 3 |
| Supplemental Figure S3 | 4 |
| Supplemental Figure S4 | 5 |
| Supplemental Figure S5 | 6 |
| Supplemental Figure S6 | 7 |
| Supplemental Figure S7 | 8 |
| Supplemental Figure S8 | 9 |
| Supplemental Figure S9 | 10 |
| Supplemental Figure S10 | 11 |
| Supplemental Figure S11 | 12 |
| Supplemental Figure S12 | 13 |
| Supplemental Figure S13 | 14 |
| Supplemental Figure S14 | 15 |
| Supplemental Figure S15 | 16 |
| Supplemental Table S1 | 17 |
| Supplemental Table S2 | 18 |
| Supplemental Method S1 | 19 |
| Supplemental Method S2 | 20 |
| Supplemental References | 21 |

**Supplemental Figure S1**. Chromatin local to *RPS7A*, encoding a small ribosomal subunit protein, changes in accordance with its repression in transcription. **(A)** Typhoon plot shows that the nucleosomes in the gene body of *RPS7A* become well-organized over the 120 min time course. **(B)** Cross-correlation heatmap shows the upstream shift of the nucleosomes in the the gene body of *RPS7A*, as well as a replacement of small fragments with nucleosome-sized fragments 200 bp upstream of its TSS. **(C)** Line plot shows that both reduced promoter occupancy and an organization of gene body nucleosomes coincide with decreased transcription of *RPS7A*.

**Supplemental Figure S2**. *CKB1*, coding for a cell growth protein kinase, exhibits unchanging transcript levels and chromatin. **(A)** Typhoon plot of *CKB1* shows transcription and chromatin do not change over the duration of the 120 min time course. **(B)** Cross-correlation heatmap shows the chromatin local to the TSS of *CKB1* is essentially constant. **(C)** Line plot showing the nearly unchanging promoter occupancy, nucleosome disorganization, and transcription of *CKB1* through 120 min.

**Supplemental Figure S3**. Changes in orthogonal chromatin measures are correlated with $\log_2$ fold-changes in transcription. **(A)** Scatter plot shows our combined chromatin score is correlated with average $\log_2$ fold-change in transcription with $r$=0.68. **(B)** Scatter plot shows average change in small fragment occupancy and average change in nucleosome disorganization are less correlated, with $r$=0.33, indicative of the orthogonal statistical power of each in correlating with changes in transcription.

**Supplemental Figure S4**. Sulfur pathway gene *MET32* activates with binding dynamics local to Met4 complex binding. **(A)** *MET32* typhoon plot shows changes in small fragments at motifs for TFs in the Met4 complex. Small fragments at a Cbf1 motif (red triangle) change from a clear well-positioned cluster at 0 min to an enrichment of small fragments near a Met4 motif (green triangle) and a Met32 motif (obscured by green triangle) from 7.5 min onward. **(B)** Cross-correlation heatmap of the dramatic chromatin changes at *MET32* beginning at 7.5 min. Changes include nucleosome disorganization, downstream shift of the +1 nucleosome, and increased occupancy of small fragments upstream of *MET32*. **(C)** Line plot shows the transcriptional activation of *MET32* at 7.5 min alongside significant nucleosome disorganization.

**Supplemental Figure S5**. Sense transcription of *MET31* is repressed while antisense transcription is induced. **(A)** Typhoon plot of *MET31* shows activation of antisense transcription within the transcript boundary of *MET31*, downstream of small fragment enrichment at a Met31/Met32 binding motif (yellow triangle). **(B)** Cross-correlation heatmap of *MET31* shows the nucleosome disorganization associated with an induction of antisense transcription. **(C)** Line plot shows the nucleosome disorganization of *MET31* is anti-correlated with its repressed sense transcription.

**Supplemental Figure S6**. Sulfur-saving *PDC6* is highly activated with dramatic changes in its local chromatin. **(A)** Typhoon plot of *PDC6* shows significant enrichment of small fragments at binding motifs for Met32 (red triangle) and Met4 (yellow triangle) beginning at 30 min. **(B)** Cross-correlation heatmap of *PDC6* shows a clear downstream shift of the +1 nucleosome and dramatic disorganization of other nucleosomes in the gene body between 30–120 min. **(C)** Line plot shows the highly activated transcription of *PDC6* is associated with significant enrichment of small fragments in the promoter and nucleosome disorganization in the gene body.

**Supplemental Figure S7**. The expression of *MCD4*, encoding an endoplasmic reticulum membrane protein, is up-regulated with evidence for changes in binding by known regulators Zap1 and Snf1 (Lyons *et al.* 2000; Venters *et al.* 2011), but also exhibits surprisingly increased nucleosome organization within its gene body. **(A)** Typhoon plot depicts the complex chromatin and transcription dynamics of *MCD4*. Along with increased transcription within the gene body of *MCD4*, a small transcript 200–300 bp upstream of the TSS is decreasing concomitantly. Time course shows monotonically changing binding at TF motifs flanking this upstream transcript. Small fragments become enriched at a Zap1 motif (yellow triangle). Small fragments at a Snf1 motif (red triangle) are replaced by fragments greater than 190 bp. **(B)** Cross-correlation heatmap depicts an enrichment of small fragments just upstream of *MCD4* as well as increased nucleosome organization. **(C)** Line plot shows an anti-correlated relationship between the decreased nucleosome disorganization and increased transcription of *MCD4*.

**Supplemental Figure S8**. *UTR2*, whose overexpression has been linked to endoplasmic reticulum stress (Miller *et al.* 2010), exhibits decreased sense transcription with increased antisense transcription. **(A)** Typhoon plot of *UTR2* shows an inverse change in transcript level on the sense and antisense strands near the 5' end of the ORF of *UTR2*. As *UTR2* is being down-regulated, an antisense transcript is becoming highly expressed. **(B)** Cross-correlation heatmap shows an organization of the +1 nucleosome of *UTR2* as well as an upstream shift of its +1 and +2 nucleosomes (though this would a downstream shift relative to the antisense transcript). **(C)** Line plot of the time course of *UTR2* depicts decreasing chromatin dynamic scores alongside decreased rates of sense transcription.

**Supplemental Figure S9**. *YBR241C*, a gene coding for a vacuole localization protein (Wiederhold *et al.* 2009), exhibits simultaneous activation of both sense and antisense transcription. **(A)** Typhoon plot of *YBR241C* shows activation of both sense transcription and antisense transcripts near the 5' transcript boundary of *YBR241C*. **(B)** Cross-correlation heatmap of chromatin local to its TSS. Gene body nucleosomes disorganize beginning at 30 min and a small factor appears to bind between the +1 and +2 nucleosomes. **(C)** Line plot of the change in chromatin and sense transcription of *YBR241C* through the time course. Its chromatin measures change more subtly than its increased sense transcription, perhaps owing to the effects of increased antisense transcription.

**Supplemental Figure S10**. Genes with evidence of marked Aft1/Aft2 binding dynamics. **(A)** Gene *LEE1*, coding for an unknown zinc-finger protein, activates with a highly enriched signal of small factor binding at an Aft1/Aft2 binding motif. **(B)** Gene *ENB1*, coding for a ferric enterobactin transmembrane transporter (Heymann *et al.* 2000), activates concurrently with a small fragment enrichment at an Aft1/Aft2 binding motif. Biochemical assays have previously identified Aft1 as a regulator for *ENB1* (Emerson *et al.* 2002).

**Supplemental Figure S11**. Precise determination of +1, +2, and +3 nucleosome positions allows for the characterization of positional changes associated with transcription. **(A)** Annotated typhoon plot of the +1, +2, and +3 nucleosomes of *RPS7A* (red lines indicate center positions over time). Nucleosomes shift upstream concomitantly with the reduction in transcription of *RPS7A*. **(B)** Heatmap of the cross-correlation of the nucleosomes and small factors local to *RPS7A* annotated with called +1, +2, and +3 nucleosome positions (red lines). Peak calling of cross-correlation values allows for tracking of positional movement of nucleosomes across time points. **(C)** Scatter plot of the cumulative positional shift of +1 and +2 nucleosomes through the 120 min time course, for all genes where the positions of those two nucleosomes could be called. A majority of these genes exhibit some downstream shift in both their +1 and +2 nucleosomes (upper right quadrant).

**Supplemental Figure S12**. Transcript boundaries are called for genes exhibiting significant transcription on their antisense strand. **(A)** Depiction of transcript boundary calling using *MET31* as an example. We call boundaries using the sum of the RNA-seq pileup across the entire time course; adding up the signal over time helps to reduce noise and offers a better chance of identifying valid antisense transcript boundaries. **(B)** Histogram of the length of called antisense transcripts. We found 667 genes that contained antisense transcripts, most of which were between 500–3000 bp long.

**Supplemental Figure S13**. Genes selected for analysis must satisfy certain criteria related to minimum ORF length and MNase-seq coverage **(A)** Distribution of ORF lengths. Genes with ORFs shorter than 500 bp (red line) were removed from further analysis. **(B)** Distribution of MNase-seq coverage within 2000 bp window around gene TSSs. Genes with fewer than 85% (red line) covered positions were removed from further analysis.

**Supplemental Figure S14**. Fragment length distributions for the first **(A)** and second **(B)** MNase replicates. The distributions of fragment lengths, for both nucleosomal lengths and shorter lengths, remain quite consistent across all time points in each replicate. **(C)** The fragment length distribution after subsampling and merging the two MNase replicates. In the merged data set, nucleosome-length fragments peak at 159$\pm$1 bp at all time points.

**Supplemental Figure S15**. Cross-correlation and entropy can be used to characterize the organizational structure of nucleosomes. **(A)** Heatmap of the nucleosome kernel computed using 2,500 well-positioned nucleosomes from Brogaard *et al.* (2012). Well-positioned nucleosome fragments are primarily between 150–170 bp in length. **(B)** Heatmap of the small factor kernel, computed using 151 Abf1 sites from MacIsaac *et al.* (2006). The kernel focuses on fragments 60–90 bp long. **(C)** Heatmap of triple nucleosome kernel, computed to quantify gene body nucleosome organization. **(D)** Typhoon plot exhibits higher values of cross-correlation at positions where MNase-seq fragments best match with nucleosome kernel (turquoise) or small factor kernel (orange). **(E)** Sum of the nucleosome cross-correlation scores at each gene's TSS shows the relative spacing between +1, +2, and +3 nucleosomes. **(F)** Example typhoon plots with low and high entropy values. Low entropy is observed with well-positioned nucleosomes. High entropy is observed when fragments are dispersed with little structure.

**Supplemental Table S1**. Gene Ontology (GO) terms for genes with the greatest organization in chromatin or decrease in transcript level. Score is the $-\log_{10}$ FDR of the term. Most of the highest scoring GO terms identified by RNA, relating to translation, are also identified by GO using changes in the chromatin.

**Supplemental Table S2**. Gene Ontology (GO) terms for genes with the greatest disorganization in chromatin or increase in transcript level. Score is the $-\log_{10}$ FDR of the term. Many of the highest scoring GO terms identified by RNA, such as those relating to sulfur assimilation, stress, and protein folding, are also identified by GO using changes in the chromatin, although some relating to sugar metabolism are identified by RNA but not by chromatin measures.

**Supplemental Method S1.** Merging replicate MNase and RNA samples

Replicates $A$ and $B$ are defined as sets of reads from six samples across the time course, while $T$ is the set of times at which those samples were taken:

$$
\begin{aligned}
A &= \{a_0, a_{7.5}, a_{15}, a_{30}, a_{60}, a_{120}\} \\
B &= \{b_0, b_{7.5}, b_{15}, b_{30}, b_{60}, b_{120}\} \\
T &= \{0, 7.5, 15, 30, 60, 120\}
\end{aligned}
$$

Separately for each replicate, the time point with the fewest reads determined that replicate's subsampling depth ($k_A$ and $k_B$):

$$k_A = \min_{t \in T} |a_t| \ , \ k_B = \min_{t \in T} |b_t|$$

The reads at each time point in each replicate were then subsampled (uniformly at random) to that replicate's subsampling depth in order to form new sets $A'$ and $B'$:

$$
\begin{aligned}
A' &= \{a'_0, \ldots, a'_{120}\} \\
B' &= \{b'_0, \ldots, b'_{120}\} \\
|a'_t| = k_A \ &, \ |b'_t| = k_B \ , \ \forall t \in T
\end{aligned}
$$

Finally, the sets of subsampled reads were merged into a superset, $M$, for downstream analysis:

$$M = \{m_0, \ldots, m_{120}\} = \{a'_0 \cup b'_0, \ldots, a'_{120} \cup b'_{120}\}$$

This procedure yields the highest possible overall read depth for the merged superset while still maintaining a consistent depth and a consistent proportion of reads between replicates $A$ and $B$ across all of the time points.

**Supplemental Method S2.** Estimating transcription rates

Transcription rates were computed for each gene using a zero-order growth with first-order decay relationship:

$$
\begin{aligned}
\frac{dC_i}{dt} &= R_i - k \cdot C_i \\
C_i &= \frac{R_i}{k} + G_i \cdot e^{-kt_i} \\
t_i &\in \{7.5, 15, 30, 60, 120\}, \text{ s.t. } i \in \{1, \ldots, 5\}
\end{aligned}
$$

where $C_i$ is the gene's total RNA concentration measured by RNA-seq for sample $i$, $k$ is its (fixed) decay rate, $G_i$ is the concentration of its RNA governed by zero-order growth, and $R_i$ is its unknown transcription rate. Assuming a constant rate of transcription between time points, we can solve for $R_i$ and $G_i$ using pairs of difference equations:

$$
\begin{aligned}
C_{i-1} &= \frac{R_i}{k} + G_i \cdot e^{-k \cdot t_{i-1}} \\
C_i &= \frac{R_i}{k} + G_i \cdot e^{-k \cdot t_i}
\end{aligned}
$$

Similarly, steady-state transcription rates, $R_0$ at $t_0$, were computed by setting the rate of production equal to the rate of decay:

$$
\begin{aligned}
t_0 &= 0, i = 0 \\
R_0 &= k \cdot C_0
\end{aligned}
$$

To compute the decay rate $k$ for a given gene, a half-life value $\tau$ was computed as an average of the values reported in three previous studies (Geisberg *et al.* 2014; Miller *et al.* 2011; Presnyak *et al.* 2015). The gene's decay rate was then set to the inverse of the average half-life, $k = 1/\tau$.

Because the above method may compute a transcription rate as being less than or equal to 0 at some time point, rates were floored to be at least 0.1 TPM/min so as to facilitate reasonable evaluations of fold change.
